# Interaction between PSD 95 and TRPV4 through PDZ domain controls TRPV4’s localization and activity

**DOI:** 10.1101/2023.06.30.547235

**Authors:** Eun Jeoung Lee, Kiwol Kim, Otgonnamjil Davaadorj, Sung Hwa Shin, Sang Sun Kang

## Abstract

The TRPV4 cation channel, is expressed in a broad range of tissues where it participates in generation of Ca^2+^ signal and/or depolarization of membrane potential. Here, we identified post synaptic density protein 95 (PSD95) as an interacting protein of this epithelial Ca^2+^ channel using confocal microscopy analysis and immunological assay. Using co-immunoprecipitation assays, we demonstrated that PSD95 was part of the TRPV4 protein complex. PSD95 protein was specifically associated with the C-terminal tail of TRPV4 to form a complex. A TRPV4 tail deletion mutant (ΔDAPL^871^: 4d) exhibited a diminished capacity to bind PSD95. Confocal microscopy analysis suggested that apical localization of TRPV4 required PSD95–TRPV4 interaction. Our data clearly suggest that formation of a complex between TRPV4 and PSD95 can regulate TRPV4 membrane localization. Both TRPV4 Ca^2+^ channel and its autophagy activity of 4d were reduced by more than 80% compared to those of the TRPV4 wild type. Our observation suggests that PSD95–TRPV4 complex plays crucial roles in routing TRPV4 to the apical plasma membrane and maintaining its authentic Ca^2+^ channel and biological function.

**Capsule:** *Background:* TRPV4 contain putative PDZ tail motif (DAPL^871^).

*Results:* Deletion of TRPV4 tail PDZ motif fails to interact with PSD95 PDZ III domain.

*Conclusion:* TRPV4 tail is an authentic PDZ motif to interact with PSD95.

*Significance:* Interaction between TRPV4 and PSD95 requires for its proper biological functions.

## Introduction

The TRPV4 cation channel, a member of the transient receptor potential (TRP) vanilloid subfamily, is expressed in a broad range of tissues where it participates in the generation of a Ca^2+^ signal and/or depolarization of membrane potential (Chun et al, 2012; Kang et al, 2012; White et al, 2016; Gao et al, 2022). The participation of TRPV4 in osmo- and mechano-transduction contributes to important functions including cellular and systemic volume homeostasis, arterial dilation, nociception, epithelial hydroelectrolyte transport, bladder voiding, and ciliary beat frequency regulation (Liedtke et al, 2007; Toft-Bertelsen et al, 2019). TRPV4 also responds to temperature, endogenous arachidonic acid metabolites, and phorbol esters including the inactive 4α-phorbol 12, 13-didecanoate (4α−PDD), and participates in receptor-operated Ca^2+^ entry, thus showing multiple modes of activation (Vriens et al, 2007; Lu et al, 2021). In this regard, several proteins have been proposed to modulate TRPV4 subcellular localization and/or function including microtubule-associated protein 7, calmodulin, with no lysine protein kinases, and PACSIN3 (Suzuki et al, 2003; Cuajungco et al, 2006; Liedtke et al, 2007; Chun et al, 2012; Kang et al, 2012; Loukin et al, 2015; White et al, 2016; Toft-Bertelsen et al, 2019; Gao et al, 2022). In addition, a close functional and physical interaction exists between the inositol triphosphate receptor 3 (IP3R3) and TRPV4 sensitizes that the latter to the mechano- and osmo-transducing messenger 5’-6’-epoxieicosatrienoic acid (EET) (Fernandes et al, 2008; Matin et al, 2016).

Post Synaptic Density 95 (PSD95) is a member of the membrane-associated guanylate kinase (MAGUK) family, which contains three PDZ domains (domain first discovered in PSD95/Dlg/ZO1 proteins), a Src-homology-3 (SH3) domain or WW motif (two conserved tryptophan residues), and a region homologous to yeast guanylate kinase (GK region) (Maruoka et al, 2000; Kim & Sheng, 2004; Magidovich et al, 2006). It has also been suggested that PSD95 may also function as a nociceptive scaffolding protein in dorsal root ganglion neurons, bringing proteins required for nociception together at the plasma membrane (Hruska-Hageman et al, 2004; Tao & Johns, 2006). PSD95 has been shown to be up regulated by inflammatory mediators, including nerve growth factor and nitric oxide. (Brenman et al, 1996; Nikonenko et al, 2008). The clustering of scaffolding molecules at contact sites is thought to play a role in the retention of specific neurotransmitter receptors at a particular synapse type (States et al, 2008; Christensen et al, 2019). A major scaffolding molecule localized at the PSD of excitatory glutamatergic synapses is the postsynaptic density protein PSD95, considered as a postsynaptic marker protein (States et al, 2008; Christensen et al, 2019).

Recently, we also demonstrated that ACE2 tail could interact with PSD95 by its putative PDZ motif tail (Lee et al, 2021; Zhang et al, 2021). PSD95 is one of the well-characterized PDZ domain containing proteins (Christensen et al, 2019; Zhang et al, 2021), and is considered as a postsynaptic marker protein (Maruoka et al, 2000; Kim & Sheng, 2004; Magidovich et al, 2006). Because PSD95 contains three PDZ domains which contribute its partner proteins’ appropriate subcellular localization and biological function, the interaction between TRPV4 and PSD95 may also control TRPV4 apical membrane localization, and its enzymatic activity, like ACE2 (Christensen et al, 2019; Lee et al, 2021; Zhang et al, 2021).

The aim of the present study was to identify proteins that interact specifically with the C-terminal tail of TRPV4 (Chun et al, 2012; Kang et al, 2012; White et al, 2016; Gao et al, 2022). In the initial screen to identify proteins associated with scaffolding proteins, PSD95 was proposed as a candidate protein because it has been reported that the TRPV4 hetero-tetramer can bind the C-terminal tail of PSD95. The functional interaction between TRPV4 and PSD95 was further substantiated by pull-down assays, confocal microscopic studies, and Ca^2+^ ion image analysis. The confocal microscopic analysis of transfected EGFP-TRPV4 WT or mutant (or ΔDAPL^871^) with PSD95 also showed that TRPV4 can interact with PSD95’s PDZ domains through its tail [DAPL^871^] motif (Chun et al, 2012; Kang et al, 2012; White et al, 2016; Gao et al, 2022). Thus, comparing with TRPV4 tail deletion mutant (ΔDAPL^871^), our data clearly identified PSD95 as a new regulatory protein that is involved in TRPV4 subcellular localization, its Ca^2+^ channel, autophagy, and apoptosis activity.

## Results

In the course of TRPV4 functional study, we recognized that it has a putative PDZ motif [DAPL-COOH^871^] in its tail (Fig 1A), which is matched with the PDZ tail type II (ΦxΦ-COOH, Φ is a hydrophobic amino acid). To find out the protein which interacts with the motif, we screened and selected Post Synaptic Density 95 (PSD95), using its consensus motif information (Fig 1A). (Garcia-Elias et al. 2008).

**Figure 1.**
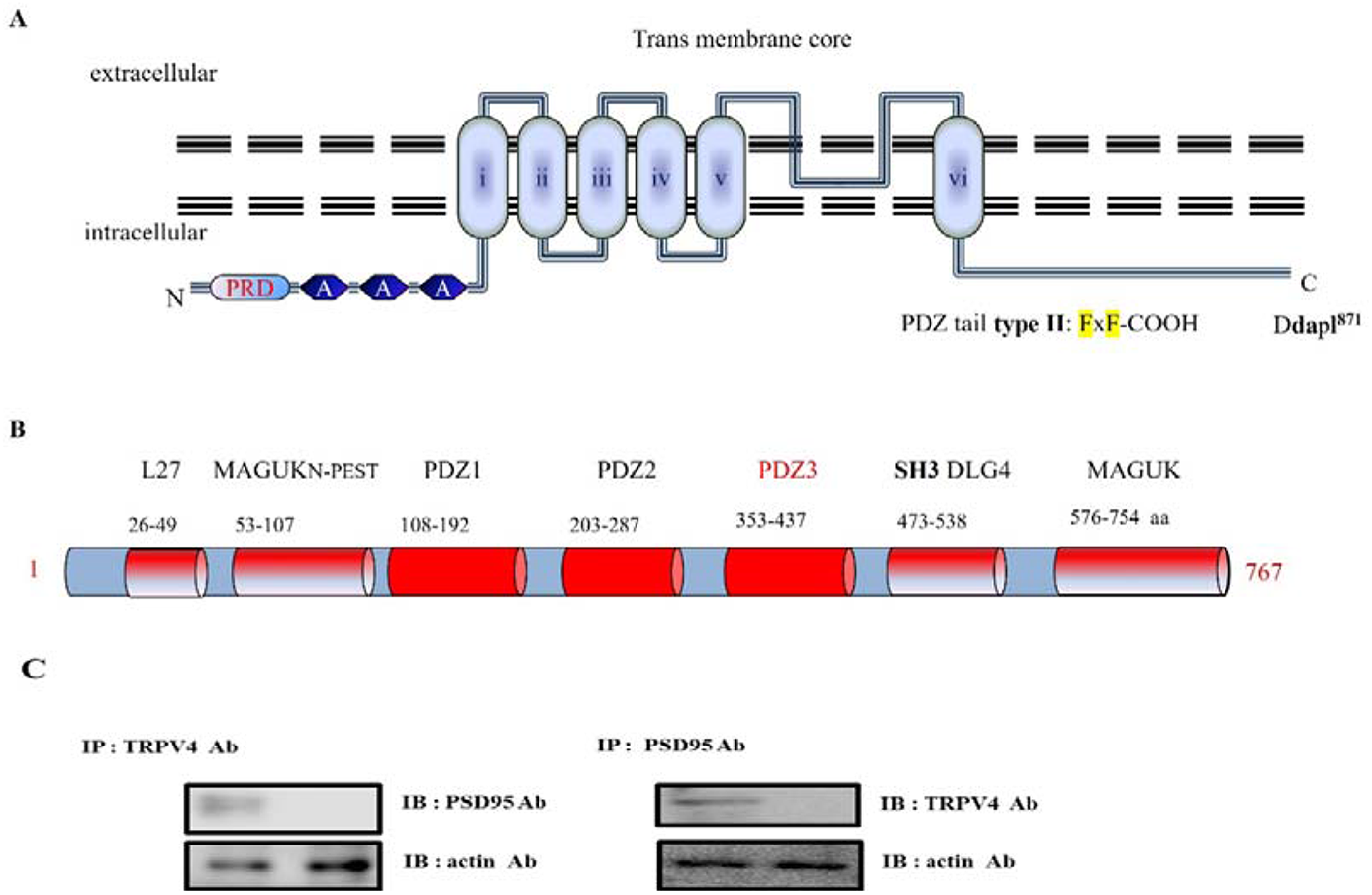
Transmembrane topology of mouse TRPV4 (871 aa) including its putative PDZ motif tail and (B) PSD95 domains. (A) Transmembrane topology of mouse TRPV4 (871 aa). The three ankyrin binding repeats (ANK; gray circles), the six trans-membrane regions (TM1_TM6), the PSD95 binding site (PSD95), and PDZ motif tail DAPL^871^are indicated. The line indicates the GST C-terminal cytoplasmic region of TRPV4 (718-871aa) or mutant (Δ827) fusion protein, the TRPV4 mutant site (Δ827-871 or ΔDAPL^871^) is compared with the wild-type (WT; Gene Bank no. BC127052). Alignment of TRPV4 WT, Δ827, and ΔDAPL^871^ with the consensus SGK1 substrate motif. Δ827 and ΔDAPL871 were constructed using site-directed mutagenesis. The result showed that TRPV4 tail (DAPL^871^) requires to interact with PSD95. (B) PSD95 (gene # AAC52113) is a member of the membrane-associated guanylate kinase (MAGUK) family, which contains 3 PDZ domains (domain first discovered in PSD95/Dlg/ZO1 proteins), a Src-homology-3 (SH3) domain or WW motif (two conserved tryptophan residues), and a region homologous to yeast guanylate kinase (GK region). So PSD95 is well known a scaffolding and a post synaptic cell marker protein. Although it is clear that the PSD-95 family of MAGUKs is critical for the maintenance of normal nicotinic synapse function, the roles of individual members have not yet been defined. The amino acid # is indicated under each corresponding domain. (MAGUK_N-PEST: Polyubiquitination (PEST) N-terminal domain of MAGUK_, SH3 DLG4: Src Homology 3 domain of Disks Large homolog 4). PSD95 Cysteine residues (3 and/or 5) are necessary for its palmitoylation (Jeyifous et al, 2016). (C) Protein-protein interaction between TRPV4 and PSD95 in HEK 293 cells. Following immunoprecipitation (IP) using an anti-TRPV4 antibody, immunoblot (IB) analysis was performed using an antibody against PSD95 (left). Conversely, PSD95 immunoprecipitated complexes were subjected to immunoblot analysis using an anti-TRPV4 antibody (right). Co-immunoprecipitation of PSD95 with TRPV4 confirmed the formation of a TRPV4–PSD95 complex. The negative control for immunoprecipitation was an unrelated antibody. The control for western blot analysis was an antibody against actin (bottom).

PSD95 was initially identified as a glutamate receptor and its cytoplasmic interacting protein in the process of excitatory synaptic transmission in the mammalian central nervous system. PSD95(#AAC52113) is a member of the membrane-associated guanylate kinase (MAGUK) family, which contains three PDZ domains (domain first discovered in PSD95/Dlg/ZO1 proteins), a Src-homology-3 (SH3) domain or WW motif (two conserved tryptophan residues), and a region homologous to yeast guanylate kinase (GK region; Fig 1B). Each functional domain and the corresponding amino acid numbers are indicated in Fig 1B. For those reasons, PSD95 is also well known as a post synaptic cell marker protein (Wyszynski et al, 1998; Keith & El-Husseini, 2008).

TRPV4-interacting protein by a putative motif search screening, and the binding between both proteins were subsequently confirmed by GST pull-down assays. The binding was not restricted to TRPV4 since TRPV4 was also shown to bind with PSD95, indicating a mutual mechanism in the regulation of these Ca^2+^ channels. Considering the high degree of homology and the similarities in electrophysiological behavior between both channels, it is indeed likely that PSD95 and TRPV4 have common regulatory factors such as associated regulatory proteins. To confirm this hypothesis, we could show by co-immunoprecipitations (Fig 1C) and GST pull-down experiments that the membrane-associated and microfilament-binding protein, PSD95, is part of the channel–PSD95 complex. Employing lysates from cells co-expressing TRPV4, we were not able to immunoprecipitate PSD95 together with TRPV4 using anti-TRPV4 antibodies (Fig 1C). The physical association of theTRPV4–PSD95 complex with TRPV4 could be too weak to resist detergent solubilization, or it could be spatially restricted to the plasma membrane, only locally regulating TRPV4 and PSD95 (Fig 1C). Because TRPV4 forms homo- and heterotetramic channel complexes, it possibly creates four PSD95 binding sites per functional channel.

### The protein-protein interaction between TRPV4 PDZ motif [DAPL-COOH^871^] and PSD95 PDZ III domain (Fig 2)

To further establish the interaction between TRPV4 and PSD95, GST pull-down binding assays were performed. As shown Fig 2A, EGFP tagging TRPV4 WT or the deletion mutant (Δ 872-871 or Δ DAPL^871^) was transfected to HEK293 cell, and each cell lysate was immunoprecitated (IP) with EFGP Ab. The immunoprecipitant was subjected to the immunoblotting (IB) with TRPV4, PSD95, or actin Ab. Even though TRPV4 WT brought down PSD95, but both deletion mutants (Δ872-871 and ΔDAPL^871^) did not bring down PSD95 (Fig 2A). The result showed that TRPV4 tail motif is enough to bind with PSD95 in HEK293 cell.

**Figure 2.**
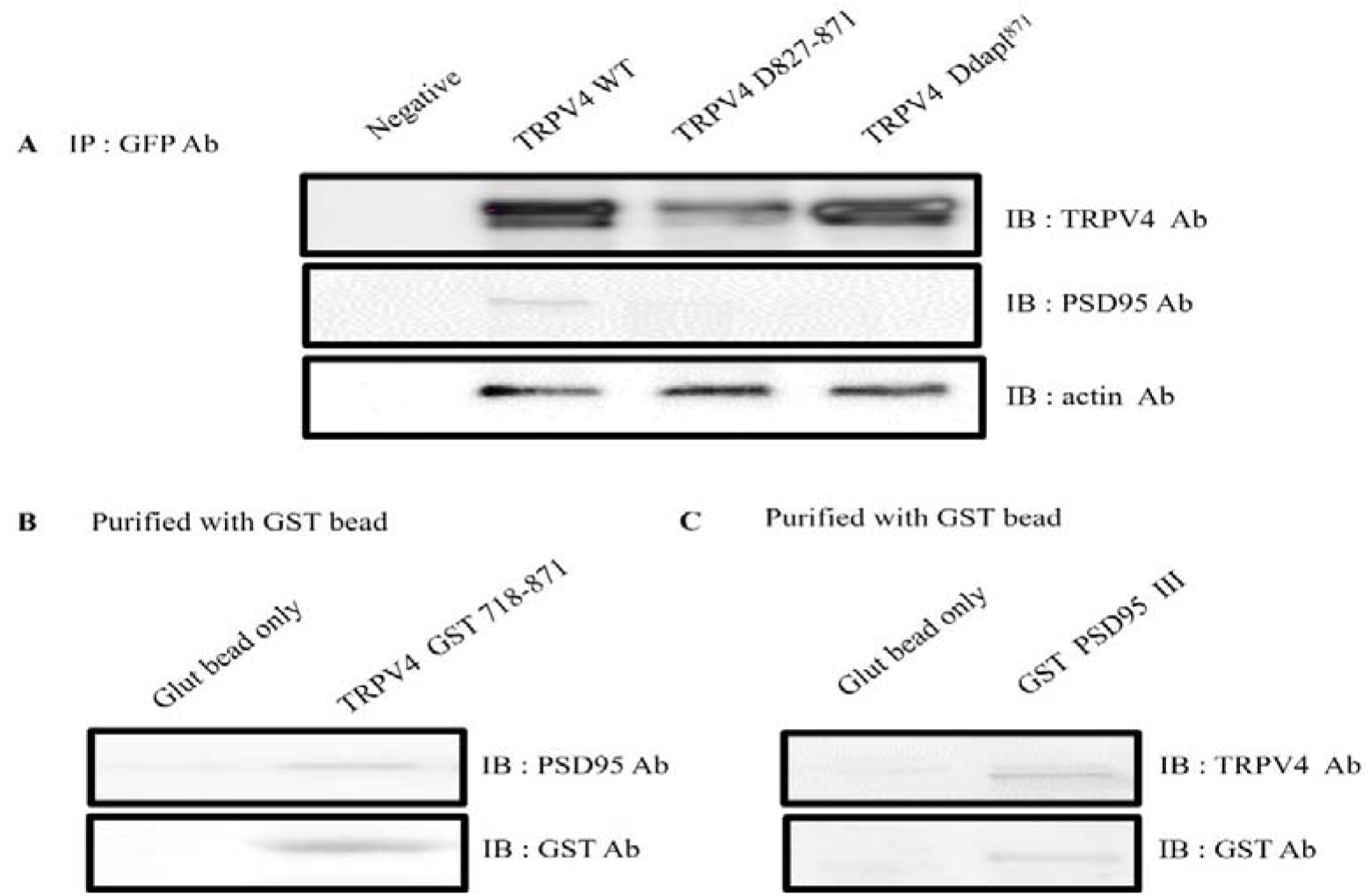
The C-terminal of TRPV4 tail (DAPL^871^) is required for interaction with PSD95. A) EGFP tagging TRPV4 WT or the deletion mutant (Δ 872-871 or Δ DAPL^871^) was transfected to HEK293 cell, and each cell lysate was immunoprecitated (IP) with EFGP Ab. The immunoprecipitant was subjected to the immunoblotting with TRPV4, PSD95, or actin Ab. Both deletion mutants (Δ872-871 and ΔDAPL^871^) did not bring down PSD95. Actin Ab western blot was used to control the equal protein amount. B) After incubation of GST-TRPV4 (718-871) with HEK293 cell lysate, the purified GST bead was immunoblotted with an antibody against PSD95 (upper lane). The GST TRPV4 the C-terminal (aa 718-871) did pull down PSD95 (right lane), whereas GST bead alone did not (left lane). C) Reciprocally, we tested whether GST PSD95 III also bring TRPV4. After incubation of GST-PSD95 PDZ III-(353-437aa) or GST bead alone with HEK293 cell lysate, the purified GST bead was immunoblotted with an antibody against TRPV4 (upper lane). The only PSD95 PDZ III did pull down TRPV4 (right lane), whereas GST bead alone did not (left lane). *In vitro*, these observations fully supported that TRPV4 tail (DAPL^871^) is enough to interact with PDZ III domain of PSD95 (Fig 1B).

To confirm it more, the pull-down assay was performed with GST -TRPV4 (718-871) or GST PSD95 PDZ III which was expressed in *E. coli*. After incubation of purified GST-TRPV4 (718-871) with HEK293 cell lysate, the purified GST bead was immunoblotted with an antibody against PSD95 (upper lane). As shown in Fig 2B, the GST TRPV4 the C-terminal (aa 718-871) was enough to pull down PSD95 (right lane), whereas GST bead alone did not (left lane).

Reciprocally, we tested whether GST PSD95 PDZ III also bring TRPV4 *in vitro*. After incubation of GST-PSD95 PDZ III-(353-437aa) or GST bead alone with HEK293 cell lysate (Cho et al, 1992), we observed that the purified GST bead was immunoblotted with an antibody against TRPV4 (Fig 2C left lane). Because the only PSD95 PDZ III did pull down TRPV4 (Fig 2C right lane), the result supported our hypothesis that TRPV4 tail (DAPL^871^) is enough to interact with PDZ III domain of PSD95 (Fig 1B).

Taken together, we concluded that the C-terminal of TRPV4 tail (DAPL^871^) is required for interaction with PSD95 PDZ III domain (Fig 1 and 2).

### PSD95 controls TRPV4 apical membrane localization through its PDZ domain interaction (Fig 3)

PSD95 is predominantly present as a heterotetrameric complex with TRPC1, which has been implicated in numerous biological processes including endocytosis, exocytosis and membrane–cytoskeleton interactions (Maruoka et al, 2000; Hruska-Hageman et al, 2004; Kim & Sheng, 2004; Magidovich et al, 2006). PSD95 is postulated to bind to the cytoplasmic face of membrane rafts to stabilize these domains, thereby providing a link to the actin cytoskeleton. The present study demonstrated the presence of PSD95 in the PSD95–channel complex, indicating a possible function in regulating channel localization and/or activity. In line with the apical membrane localization of PSD95 and its postulated function in organizing certain plasma membrane domains, our findings provided the first functional evidence for a regulatory role of PSD95 controlling Ca^2+^ channel trafficking. Pearson’s correlation coefficient (PCC) between TRPV4 WT and PSD95 was 0.89 +/-0.04 (n=5), whereas PCC between TRPV4 ΔDAPL^871^and PSD95 was 0.07 +/-0.05 (n=5) in the confocal microscopy (Fig 3). Consistent with the Western blot result in Fig. 1 and 2, the confocal result (Fig 3) also confirmed that the protein-protein interaction between TRPV4 and PSD95 through the tail motif (DAPL^871^) of TRPV4 is required for its apical membrane localization with PSD95.

**Figure 3.**
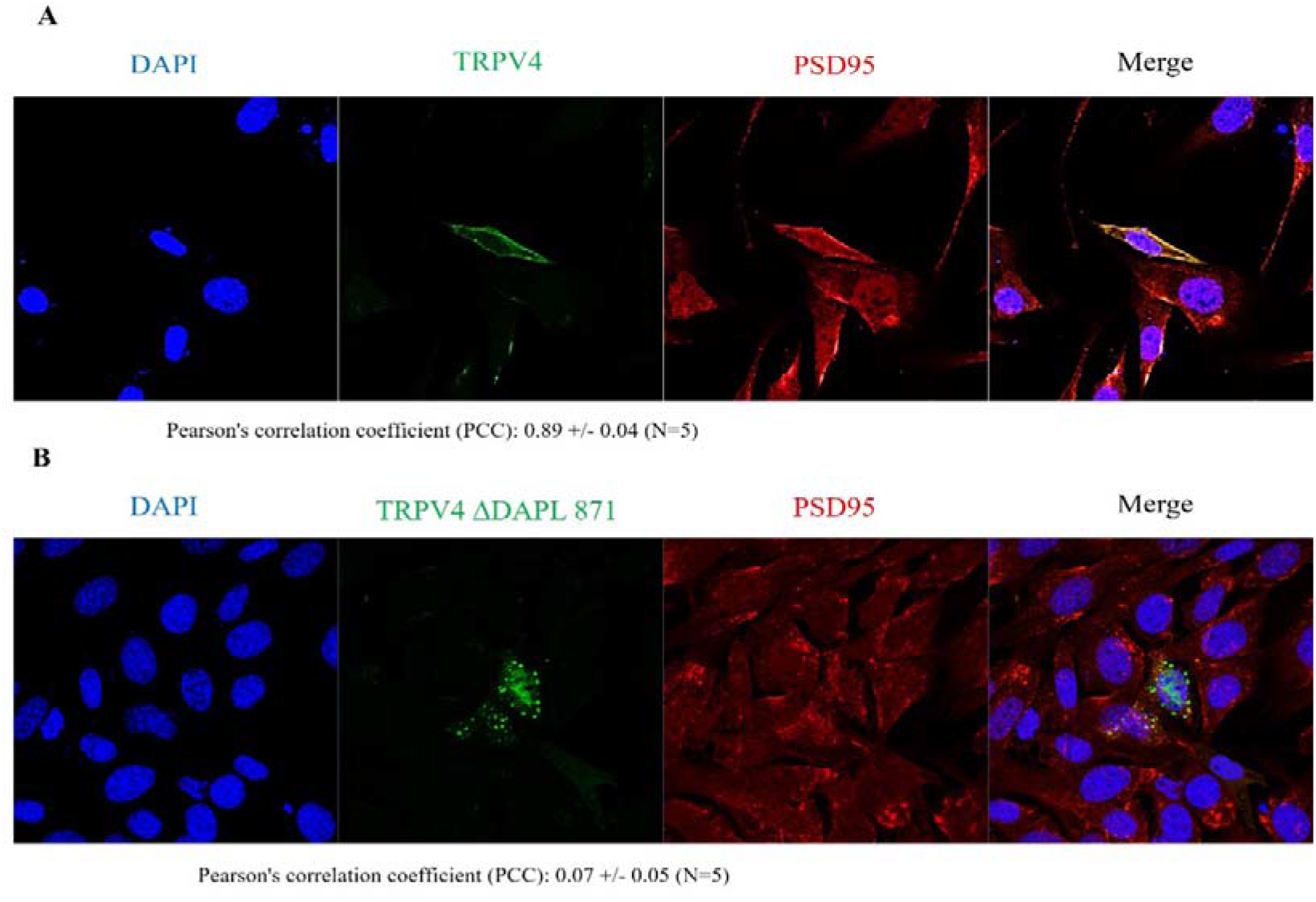
Confocal microscopic images of cells transfected with EGFP-TRPV4 WT, or ΔDAPL^871^. Cells were examined by direct immunofluorescence microscopy. The figures show EGFP-TRPV4 WT or mutant (ΔDAPL^871^) (green), PSD95 (red), and merged (yellow) confocal microscopic images. EGFP-TRPV4 WT or DDAPL871 showed minimal co-localization with PSD95 at the Golgi apparatus (B left and right). EGFP-TRPV4 DDAPL^871^ was principally detected in the plasma membrane (right). However, EGFP-TRPV4 WT was co-localized with PSD95 at the Golgi apparatus (B middle). Fn/c (factional rate of nuclear localization to cytoplasm and PCC (Pearson Co-localization co-efficiency) are indicated below.

### The proper TRPV4 subcellular localization is necessary for its Ca^2+^ channel activity (Fig 4)

The effect of PSD95 on TRPV4 and TRPV4 activity was determined by whole-cell patch–clamp analysis in transiently transfected human embryonic kidney (HEK293) cells. As shown in Fig 4 (one of five repeats), Ca^2+^ activity of 4d (ΔDAPL^871^: lower graph) was reduced by more than 80% compared to that of TRPV4 wild type (upper graph) in present with Fura-2 and 4 α−PDD. The un-transfected human embryonic kidney (HEK293) cells did not show TRPV4 Ca^2+^ channel activity. Thus, this observation strongly suggested that the protein-protein interaction between PSD95 PDZ III domain and TRPV4 tail motif [DAPL^871^] is necessary for TRPV4 Ca^2+^ channel activity regulation.

**Figure 4.**
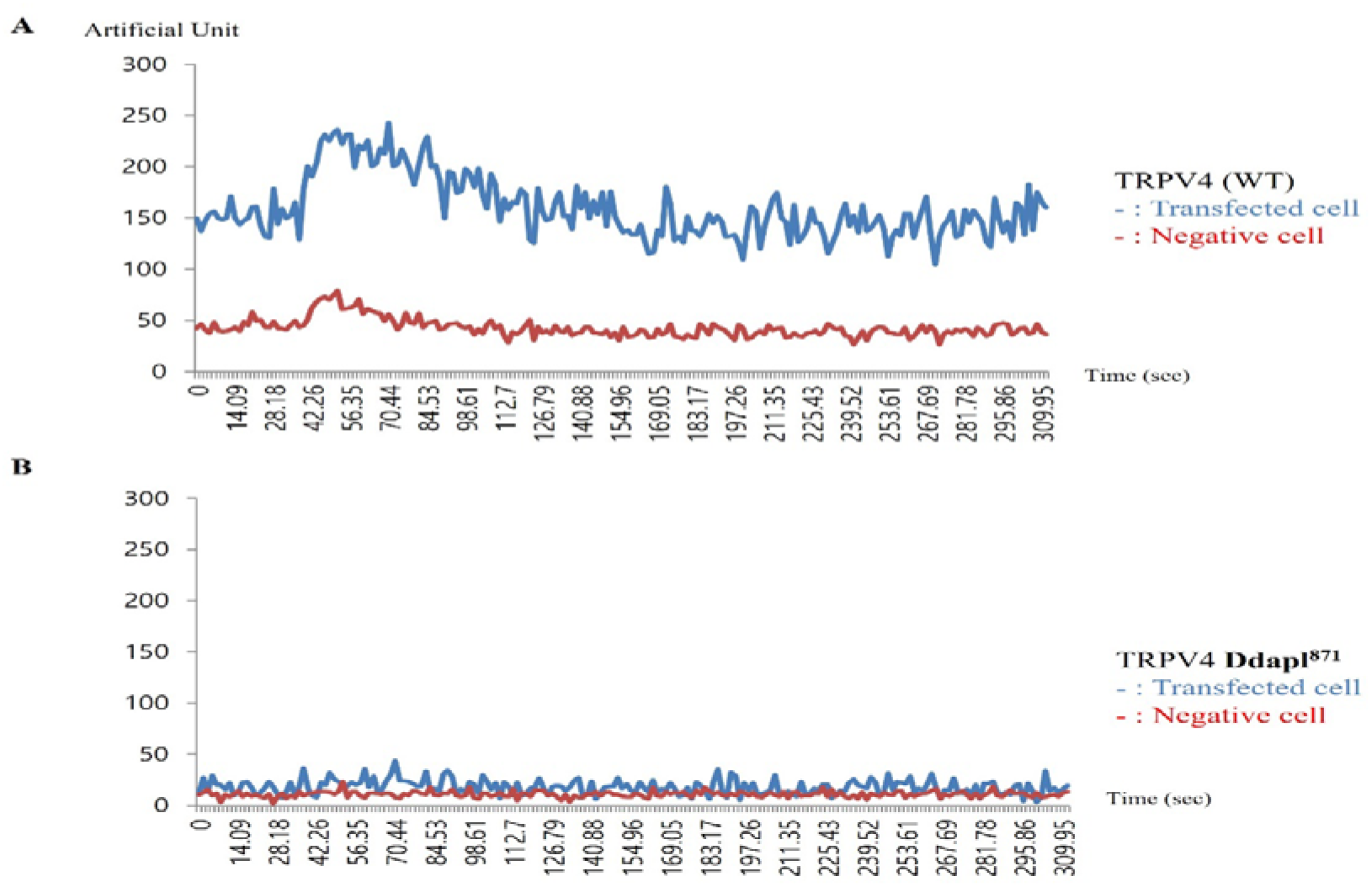
Ca^2+^ live image Comparison of TRPV4 WT, or ΔDAPL^871^ in HEK cells. Effects of 4-α PDD on intracellular calcium concentration change [Ca^2+^]I of TRPV4 WT (A), or ΔDAPL^871^(B), expressed by the absorption at 488 nm of argon-ion laser in HEK 293 cells (as an arbitrary % unit). The Ca^2+^ activity of 4d (ΔDAPL^871^: lower graph) were reduced more than 80% those of TRPV4 wild type (upper graph) in present with Fura-2 and 4 α−PDD. The image is representative of five repeat experiments.

### PSD95 also controls both TRPV4 autophagy and apoptosis activity through its PDZ domain interaction (Fig 5)

Because the d4 mutant of TRPV4 failed to produce significant Ca^2+^ currents in HEK293 (Fig 4), even though the channel was readily detectable by immunoblotting, we wonder whether the d4 mutant of TRPV4 shows the different autophagy activity or apoptosis ability, compared to those of TRPV4 WT.

**Figure 5.**
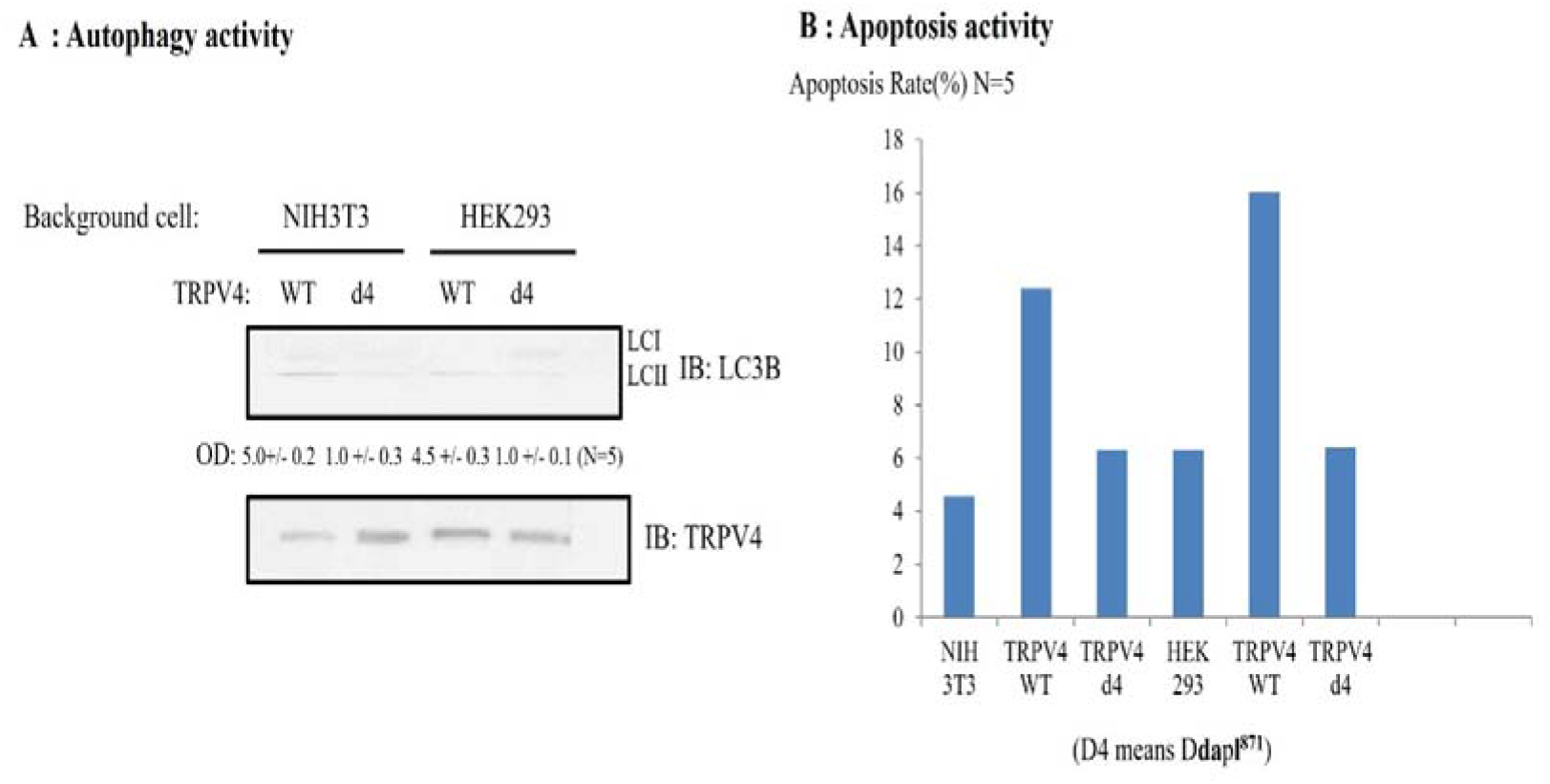
The effect the interaction between TRPV4 and PSD95 on TRPV4 autophagy and apoptosis activity. A) Comparison between TRPV4wt and d4(ΔDAPL^871^) autophagy activity. The accumulation of microtubule-associated protein light chain 3-II (LC3-II) in cells was indicated with the arrow. The same number of NIH3T3/ HEK293 cells was infected with TRPV4 WT or d4 for 48 h, then LC3-II appearance was analyzed by Western blotting (upper lane). The amount of TRPV4 protein in the experiment was monitored by TRPV4 antibody (lower lane). The relative optical density (OD) was measured by image analysis of the dried SDS-PAGE gel with the Fuji Image Quant software (Fujifilm, Tokyo, Japan), according to the manufacturer’s instructions. The number of LC3 II OD under the picture was the average of five repeats. The antibody against LC3 was used as manufacture’ recommendation. In both HEK 293 and 3T3 cell, TRPV4 mut ant (d4) autophagy activity was reduced by 80% than those of TRPV4 WT. B) EGFP-TRPV4 WT, its d4 mutant (ΔDAPL^871^) or EGFP vector was transfected and the rate of apoptosis measured by FACS. TRPV4 mutant (d4) did not promoted cell apoptosis significantly, compared to TRPV4 WT, suggesting that this is an apoptotic autophagy. For details, see the result section.

To determine whether TRPV4 autophagy activity or apoptosis ability is changed by its tail deletion, TRPV4 autophagy activity or apoptosis ability were measured with western blot or FACS (Fig 5). For comparison between TRPV4wt and d4 (ΔDAPL^871^) autophagy activity, the accumulation of microtubule-associated protein light chain 3-II (LC3-II) in cells was indicated with the arrow. The same number of NIH3T3/ HEK293 cells was infected with TRPV4 WT or d4 for 48 h, then LC3-II appearance was analyzed by Western blotting (upper lane). The amount of TRPV4 protein in the experiment was monitored by TRPV4 antibody (lower lane). The relative optical density (OD) was measured by image analysis of the dried SDS-PAGE gel with the Fuji Image Quant software (Fujifilm, Tokyo, Japan), according to the manufacturer’s instructions (Klionsky et al, 2021). The number of LC3 II OD under the picture was the average of five repeats. The antibody against LC3 was used as manufacture’s recommendation (Klionsky et al, 2021). We observed that the 4d mutant TRPV4 autophagy activity was reduced by more than 80% compared to that of the TRPV4 wild type in Fig 5A,

To compare TRPV4 autophagy activities, we also measured the apoptosis rate either TRPV4 WT or its d4 mutant (ΔDAPL^871^) or EGFP vector. Each construct w as transfected and the rate of apoptosis was measured by FACS (Shin et al, 201 2; Lee et al, 2021). As shown in Fig 5B, the 4d mutant TRPV4 apoptosis activity of was also reduced by more than 80% compared to that of the TRPV4 wild type, suggesting that TRPV4 autophagy activity is not an apoptotic autophagy (Fig 5).

Taken together, TRPV4 tail motif [DAPL^871^] is also required for both its proper autophagy and apoptosis activity through binding to PSD95 PDZ III domain.

### Our working model of [ΔDAPL^871^] **in** Ca^2+^-channel activity of TRPV4 (Fig 6)

TRPV4 is predominantly present as a homo or heterotetrameric complex with TRPV family, which has been implicated in numerous biological processes including endocytosis, exocytosis and membrane–cytoskeleton interactions (Liedtke & Heller, 2007; Chun et al, 2012; White et al, 2016). Because PSD95 is postulated to bind to the cytoplasmic face of post synaptic cell apical membrane rafts to stabilize these domains, it provides a link to the actin cytoskeleton (Fig 6). There are many proteins to integrate the synapsis structure, including in Fig 6. The present study demonstrated the presence of PSD95 in the PSD95–channel complex, indicating a possible function in regulating TRPV4 channel in the apical region of post synaptic neuronal cell, including localization and/or activity (Cuajungco et al, 2006; Fernandes et al, 2008). Furthermore, we showed here that [ΔDAPL^871^] motif in TRPV4 regulates not only the protein-protein interaction with PSD95 directly but also its autophagy and apoptosis activity (Fig 5).

**Figure 6.**
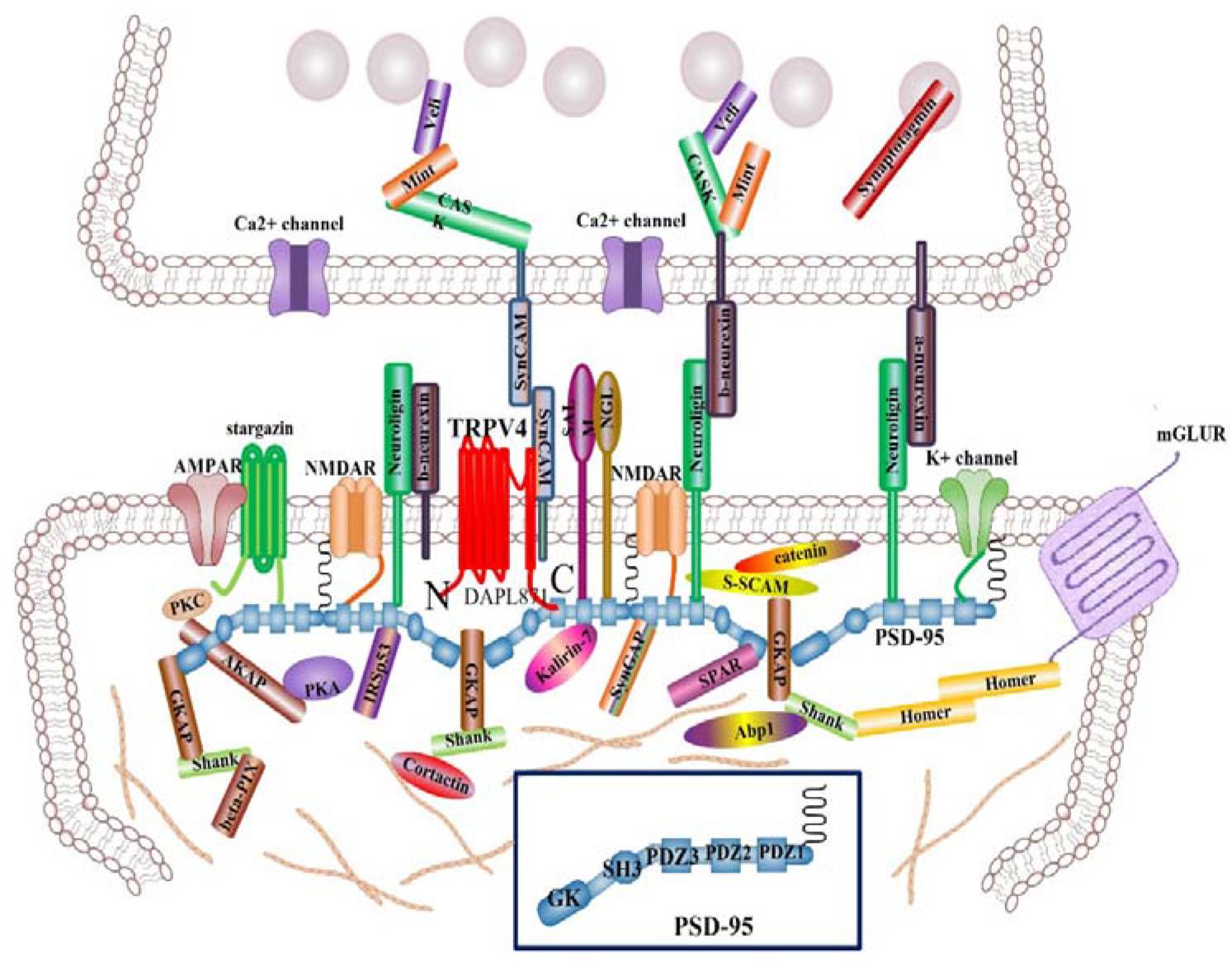
Putative model for the regulation of TRPV4 by PSD95 in the post synaptic cell. TRPV4 WT can be activated by the putative dual (activator/inhibitor) function of proteins such as PSD95 through association/dissociation from its C-terminal cytoplasmic domain and C-terminal phosphorylation. The molecular organization of postsynaptic cell is presented, but only major molecules associated with PSD95 are shown (Keith & El-Husseini, 2008; Klionsky et al, 2021). The various molecules portrayed regulate synapse function, morphology, trafficking and localization of adhesion molecules and neurotransmitter receptors. The association between PSD95 PDZ III domain and TRPV4 C-terminal PDZ motif tail (DAPL^871^) will provide an explain why TRPV4 subcellular localization is localized on the apical region of post synaptic neuron, comparing TRPV1 and 2. Interaction between TRPV4 and PSD95 may also contribute the proper postsynaptic neuron efficacy, including ACE2 (Lee et al, 2021; Zhang et al, 2021). PSD95 Cysteine residues (3 and 5) are necessary for its palmitoylation (see Fig 1B). The PDZ association between TRPV4 tail and PSD95 PDZ III domain will be one of the answers why TRPV4 is also localized in the postsynaptic neuron (Keith & El-Husseini, 2008; Shibasaki et al, 2015)

In synapsis, TRPV4 tail motif interaction with PSD95 PDZ domain may maintain TRPV4 apical subcellular localization of postsynaptic cell and contribute to synapse efficacy (Cuajungco et al, 2006; Fernandes et al, 2008; Fig 6). Even though the biological meaning of interaction between TRPV4 tail motif and PSD95 PDZ III domain in vivo should be studied further, our results firmly suggested that TRPV4 tail motif interacts with PSD95 PDZ III domain. With the PDZ motif consensus information, we noticed that TRPV3-6 contain PDZ tail type II motif, but TRPV1-2 do not (Table 1). Based on this information, we assume that TRPV3-6 also exist in the apical membrane dominantly, for their proper biological role by PSD95, similar with TRPV4. Other proteins having PDZ domain such as NHERF2, also interact with the TRPV3-6(Table 1). The further experiment results will be supported the hypothesis soon.

**Table 1.** Comparison TRPV family protein PDZ motif in its tail. It is recognized that TRPV4 (including TRPV3, 5, and 6) contains PDZ tail type II (ΦxΦ−COOH), except TRPV1 and 2. These valinoid channels can form homo/hetero tetramer. Therefore, it seems to be that PSD95 also controls the subcellular localization of TRPV1 and 2 through PDZ domain interaction, as similar with that of TRPV4 (Fig 3). TRPV4-6 contain PDZ tail type II motif, but TRPV1-2 do not. So we assume that TRPV1-6 also exist in the apical membrane dominantly, for their proper biological role by PSD95, similar with homo tetramer TRPV4. Other proteins having PDZ domain such as NHERF2 may also interact with TRPV4-6.

| PSD95 PDZ III domain binding motif (PDZ tail type II: FxF-COOH) |  |
| --- | --- |
| TRPV6 | EYQ**I**<sup>800</sup> |
| TRPV5 | VYh**F**<sup>729</sup> |
| TRPV4 | **DAPL**<sup>880</sup> |
| TRPV3 | ETs**V**<sup>791</sup> |
| TRPV2 | lqsn<sup>764</sup> |
| TRPV1 | sgek<sup>839</sup> |

In summary, we provided the first evidence of a regulatory role of the PSD95– TRPV4 complex in Ca^2+^ (re)absorption in general and in TRPV4 routing in particular. The interaction between PSD95 and TRPV4 was regulated by the phosphorylation on TRPV4. The elucidated molecular mechanism involves tethering the PSD95–TRPV4 complex to the Ca^2+^ channel (Fig 4), resulting in functional polar plasma membrane localization (Fig 3 and 6). This mechanism is likely to be applicable to other ion transporters given the wide tissue distribution of TRPV4 and the associated PSD95 protein. The TRPV4 [DAPL^871^] seems to be also the major regulatory point for the interaction with other proteins, including PSD95 and its autophagy and apoptosis activities (Fig 5).

## Discussion

The present study identified PSD95 pair as the first auxiliary protein complex for the newly identified epithelial Ca^2+^ channels, TRPV4. PSD95 forms a well-defined heteromeric complex with PSD95 and associates specifically with the conserved sequence [ΔDAPL^871^] motif located in TRPV4.

Our results show that Ca^2+^ store depletion triggers the translocation of TRPV4-C1 heteromers to the plasma membrane, resulting in an enhanced Ca^2+^ influx in response to flow. This scheme provides a mechanistic explanation for the potentiated Ca^2+^ influx to flow under the condition of Ca^2+^ store depletion in vascular endothelial cells.

Our results indicate that the [ΔDAPL-COOH^871^] mutant impairs the sensitivity to extracellular Ca^2+^ of divalent cation entry operated by the intracellular Ca^2+^ stores. Furthermore, the PSD95 PDZ III domain mutation, inducing charge swapping, which prevents PSD95-TRPV4 interaction, also results in loss of function of plasma membrane-located PSD95. These findings, together with the high Ca^2+^-selectivity of Orai1 channels in the presence of extracellular Ca^2^ ^+^, suggest that PSD95 located in the plasma membrane modulates store-operated divalent cation entry by interaction with TRPC1 channels. Despite the function of ER-located PSD95 has been widely investigated, the functional properties of plasma membrane-resident PSD95 remain largely uncharacterized.

In our results, PSD95 binds to TRPV4 by its protein interaction, not by its electrostatic charge (Fig 1 and 2). PSD95 located in the apical plasma membrane has also been reported to modulate the function of the SOC channels in different non-excitable cells (Keith & El-Husseini, 2008; Fig 6). In the present study we have further investigated the regulatory role of SOC by extracellular Ca^2+^ mediated by plasma membrane-resident PSD95 in HEK-293 or 3T3 cells, which endogenously express PSD95 and different Ca^2+^ channels, such as Orai1 and TRPC1, and allowed us to express different PSD95 mutants. Furthermore, we have attempted to characterize the mechanism involved in the modulation of store-operated divalent cation-permeable channels by plasma membrane-located PSD95. **So** PSD95 functions as an TRPV4 scaffolding protein on postsynaptic efficacy (Fig 6).

PDZ tail [ΔDAPL-COOH^871^] of TRPV4 plays a role in its apical membrane localization and forms TRPV4 cluster to facilitate COVID19 spike protein binding, resulting in enhanced infectivity of the virus. Thus, viral invasion through endocytosis with TRPV4 can be blocked by interrupting the interaction between the PDZ domain of PSD95 and its PDZ tail [ΔDAPL-COOH^871^] of TRPV4. It has been recognized that the E protein of COVID19 contains PDZ type II motif (xΦxΦ-COOH Φ, a hydrophobic amino acid) and DLLV-COOH^75^ in (^1^mysfvseetg tlivnsvllf lafvvfllvt lailtalrlc ayccnivnvs lvkpsfyvys rvknlnssrv pd**L**l**V**^75^), which seem to inhibit protein interaction by competing with its own type II PDZ tail through a feed-back inhibition mechanism (Lee et al, 2021). Thus, it seems that Type III motif (xD/ExΦ-COOH: Φ, a hydrophobic amino acid, and VDDI-COOH^85^) is able to interrupt the interaction between TRPV4 and PSD95 (Maruoka et al, 2000; Hruska-Hageman et al, 2004; Kim et al, 2004; Magidovich et al, 2006; Tao et al, 2006; Nikonenko et al, 2008). Therefore, the interaction between PDZ of TRPV4 and PSD95 seems to be the major target to intervene COVID19 related diseases. It might be one of the most potential targets for drug development. TRPV4’s tail PDZ motif [ΔDAPL-COOH^871^] can also interact with other PDZ proteins such as NHERF2 (Fig. 2). However, the other PDZ domain containing proteins to bind TRPV4’s tail PDZ motif are not clearly identified yet.

We also noticed that TRPV4 (including TRPV3, 5, and 6) contains PDZ tail type II (ΦxΦ−COOH;, except TRPV1 and 2 with its PDZ tail type II (ΦxΦ−COOH) (Table 1) . Therefore, it seems to be that PSD95 also controls the subcellular localization of TRPV3, 5 and 6 through PDZ domain interaction, as similar with that of TRPV4 (Fig 3). However, further researches are required to confirm the hypothesis. Recent research into the role of PSD95 in recruitment of glutamate receptors and adhesion molecules at the synapse has brought much insight into the basic principles governing assembly of protein complexes at glutamatergic synapses (Brenman et al, 1996; Nikonenko et al, 2008; States et al, 2008). It is evident from recent studies that scaffolding molecules such as PSD 95 play prominent roles in the assembly of ion channels and associated proteins at excitatory synaptic sites. Recent in vitro data also revealed that PSD 95 might influence the retention of particular adhesion molecules at excitatory contacts at the expense of inhibitory synapses (Fig 6). Thus it is attractive to propose that these molecules act as the molecular sensors that control the balance between excitation and inhibition.

In order to find differences between PSD95 and TRPV4, TRPV4 and PSD95 C-terminal amino acid sequences were compared among vertebrates, including human, monkey, mouse, rat, bat, chicken, snake, and xenopus. Thus, we found out that although the number of amino acids varied among species, type 1 PDZ motif [xΦxΦ-COOH] was highly conserved, suggesting that the interaction between PSD95 and TRPV4 was kept during evolution. Even though PSD95 was not limited at the apical membrane, but localized on the whole plasma membrane (Fig 6). Thus, it seems to be valuable to point out that PSD95 C-terminal amino acid sequences of mammals are much different from those of non-mammals. The interaction TRPV4 tail motif with PSD95 PDZ domains may guide TRPV4 to form a patch on the apical plasma membrane where TRPV4 can play a role membrane potential (Fig 6). Thus, disrupting the interaction of TRPV4 with PSD95 PDZ domains might be a novel method to inhibit COVID19 attachment to host cell membrane. There are many small molecules such as anti-cancer drugs that can disrupt PDZ interaction (Maruoka et al, 2000; Hruska-Hageman et al, 2004; Kim et al, 2004; Magidovich et al, 2006; Keith & El-Husseini, 2008). Therefore, anti-cancer drugs that can disrupt PDZ interaction might be useful for preventing COVID19 invasion.

Recently, others showed that PSD95 is required for thapsigargin-induced translocation of TRPV4-C1 heteromeric channels to the plasma membrane. The crucial role of Orai1 in regulating SOCE in HUVECs also has been demonstrated to suggest that PSD95 and Orai1 act through heteromeric TRPV4-C1 channels to exert its effect on SOCE (Leidtke & Heller, 2007; Vriens et al, 2007; Chun et al, 2012; Kang et al, 2012; White et al, 2016; Toft-Bertelsen et al, 2019; Gao et al, 2022). These data strongly suggested that both TRPC1 and TRPV4 play a role as SOCE in vascular endothelial cells. They also suggested that Ca^2+^ store depletion enhances trafficking of heteromeric TRPV4-C1 channels to the plasma membrane where these channels contribute to SOCE and ISOC in a PSD95- and Orai1-dependent manner. Because of these, we cannot rule out the possibility that the interaction between PSD95 and TRPV4 is mediated by other proteins, such as TRPV 3, 5, 6 and 7.

Previously, we have already demonstrated that TRPV4 cation channel is one of SGK1 authentic substrate proteins, and that the Ser 824 residue of TRPV4 is phosphorylated by SGK1 (Shin et al, 2012; White et al, 2016). In addition, we demonstrated that phosphorylation on the Ser 824 residue of TRPV4 is required for its interaction with F-actin, using TRPV4 mutants (S824D) and its proper subcellular localization (Shin et al, 2012). In the line of expansion, we also observed that the phosphorylation of the Ser824 residue promotes its dissociation from PSD95 and the plasma membrane localization (data not shown). Our results also provide evidence for a functional role of plasma membrane-resident TRPV4 in the regulation of store-operated Ca^2+^entry but, not by PDZ tail motif (ΔDAPL^871^).

Because it has been reported that PSD95 PDZ III domain binding motif PDZ tail type II, it can be bind to TRPV6 (WEYQI^800^), TRPV5 (vYhF^729^), TRPV4 (DAPL^880^) and TRPV3 (eTsV^791^). However, TRPV2 (lqsn^764^) and TRPV1 (sgek^839^) do not bind to PSD95 PDZ III directly. It is interesting to figure out the difference meaning among TRPV family proteins (Table 1). Even though TRPV2 (lqsn^764^) and TRPV1 (sgek^839^) do not bind to PSD95 PDZ III directly, TRPV1-2 are able to bind with PSD95 through forming a hetero tetramer with TRPV4. Thus, it seems to be that PSD95 controls these 6 members of TRPV channels apical membrane localization. This activity may be related with its autophagy or apoptosis activity, even though the mechanism how TRPV4 involved in this time is not clear yet (Fig 5). However, considering the existence of numerous molecules that regulate this process, future studies are needed to address mechanisms that control cross-talk between various scaffold molecules and adhesive systems to define the contribution of each system in the control of synaptic strength and brain function.

In conclusion, we provided the first firm evidence protein-protein interaction between PSD95–TRPV4 through PDZ domain and motif. The interaction between PSD95 and TRPV4 regulates its Ca^2+^ channel activity, apoptosis, autophagy, and apical membrane localization. Therefore, the further elucidation molecular mechanism of the PSD95–TRPV4 complex and identification of unknown protein in synapsis will provide more information, relating axon polarity and synapsis efficacy (Keith & El-Husseini, 2008; Fig 6).

## Materials and methods

### Site-directed mutagenesis

In order to obtain the mutants, amino acid changes were introduced using mutated oligonucleotides for 827 stop (up 5′-agg gat cgt tgg Gcc Gcg gtg gtg ccc cgc gta-3′, down 5′-gcg ggg cac cac cgC ggC cca acg atc cct acg-3′) or 867 stop (up 5′-agg gat cgt tgg GAc GAC gtg gtg ccc cgc gta-3′, down 5′-gcg ggg cac cac GTC gTC cca acg atc cct acg-3′) and wild-type TRPV4 as a template. The TRPV4 mutant constructs were prepared using a QuickChange® Multi Site-Directed a genesis Kit (Stratagene). To obtain truncated TRPV4 (aa 718–871), we used primers (up 5′-ata gga tcc atg ggt gag acc gtg ggc cag-3′, down 5′–ata ctc gag cta cag tgg ggc atc gtc cgt-3′). All TRPV4 mutants were confirmed via DNA sequencing. Human embryonic kidney (HEK293) cells were transfected with TRPV4 and with mutant constructs, as described previously.

### Glutathione S-transferase (GST)-TRPV4 fusion proteins and pull-down assays

TRPV4 sequences were PCR-amplified, subcloned into pGEX-5X-1, sequenced, and expressed in *Escherichia coli* BL21. GST-PDZ III-PSD95 (353-437aa), or GST-TRPV4 fusion proteins bound to glutathione-Sepharose were equilibrated in PBS buffer containing 0.1% Triton X-100 and 1 mM CaCl_2_ or 2 mM EGTA. Incubation with the total cell lysates or Δ827 was followed by three washes with the appropriate buffers, and the bound proteins were eluted with sample buffer, subjected to SDS gel-electrophoresis, and exposed to X-ray film (Fuji Las 3000 mini). PDZ3-PSD95 (353-437aa)-pGEX-2T expression vector was obtained from Dr. Kennedy Lab Neuron 9, 929-942 (1992) (Cho et al, 1992).

### Fluorescence measurements of [Ca^2^ ^+^]_i_

We measured [Ca^2^ ^+^]_i_ using a fluorescent Ca^2^ ^+^ indicator Fluo4-acetoxymethyl ester (Fluo4-AM), as previously described. In brief, cells growing on coverslips were incubated for 40 min in DMSO solution containing 1 μM Fluo4-AM at 24 °C in darkness, and then washed and incubated for 15 min to hydrolyze internalized Fluo4-AM. We measured [Ca^2^ ^+^]_i_ in single cells that emitted fluorescence, using confocal microscopy (LSM710 Zeiss, Germany) at wavelengths of 495 nm (excitation), and 519 nm (emission). The absorption (as an arbitrary unit) at 488 nm with an argon-ion laser was measured as a relative intracellular Ca^2^ ^+^ ion concentration [Ca^2^ ^+^]_i_. All experiments were carried out at 24 °C. After stimulation with mild heat (from 24 to 42 °C within 45 s for 2 min), [Ca^2^ ^+^]_i_ was measured in single cells at a 24 °C solution. (Shin et al, 2012).

### Solutions and drugs

Cells were superfused normally with a solution containing (mM): 88 NaCl, 5 KCl, 5.5 glucose, 1 CaCl_2_, 10 HEPES and 100 mannitol, adjusted to pH 7.4 with NaOH (300 mosm kg^−1^ H_2_O). The HTS was adjusted to 200 mosm kg^−1^ H_2_O by omitting mannitol. Fluo4-AM was acquired from Sigma (St Louis, MO, USA). Stock solutions of phorbol esters were initially prepared in dimethyl sulphoxide (DMSO) at a concentration of 1 mM, and then stored at −20 °C. The final DMSO concentration in the experimental bath solution containing phorbol esters never exceeded 0.5%.

***Autophagy assay*—**to measure autophagy, we used LC3 western blot (Bio-Protocol Com) method following the manufacture’ guide. Biochemical methods such as western blot to measure autophagic proteins have been used in autophagy studies. The western blot was performed same as the above “Immunoblotting”section with an antibody against LC3. Each assay was performed in five replicates.

### Statistics

Data are expressed as the means ± standard error of the mean (S.E.M.). Statistical significance was determined via Student’s *t*-test or Welch’s *t*-test. We utilized the Mann– Whitney test to compare the effects of 4-αPDD to normal conditions. Values of P < 0.05 were considered statistically significant.

## Acknowledgements

This work was supported by Korea Research Foundation Grant (KRF-2013-331-C00224) and Chungbuk Natural Science Foundation [Grant Numbers 7131003 and 5112007] to S. S. Kang.

